# Immune engineered extracellular vesicles to modulate T cell activation in the context of type 1 diabetes

**DOI:** 10.1101/2022.11.23.517693

**Authors:** Matthew W. Becker, Leeana D. Peters, Thinzar Myint, Todd M. Brusko, Edward A. Phelps

**Affiliations:** J. Crayton Pruitt Family Department of Biomedical Engineering, University of Florida, Gainesville, FL; Department of Pathology, Immunology, and Laboratory Medicine, College of Medicine, University of Florida, Gainesville, FL; University of Florida Diabetes Institute, University of Florida, Gainesville, FL; Department of Pediatrics, College of Medicine, University of Florida, Gainesville, FL

**Keywords:** Extracellular vesicles, immune engineering, PD-L1, CD80, type 1 diabetes

## Abstract

Extracellular vesicles (EVs) are small, biologically active, cell-secreted vesicles that can affect immune responses through antigen presentation and co-stimulation or co-inhibition. We generated designer EVs to modulate autoreactive T cells in the context of type 1 diabetes by engineering K562 cells to express HLA-A*02 (HLA-A2) alongside co-stimulatory CD80 and/or co-inhibitory PD-L1. EVs presenting HLA-A2 and CD80 activated CD8^+^ T cells in a dose, antigen, and HLA-specific manner. Adding PD-L1 to these EVs produced an immunoregulatory response, reducing CD8^+^ T cell activation and cytotoxicity *in vitro*. EVs alone could not stimulate T cells without antigen presenting cells (APCs), suggesting that EVs act by cross-dressing APCs. EVs lacking CD80 were ineffective at modulating CD8^+^ T cell activation, suggesting that both peptide-HLA complex and costimulatory molecules are required for EV-mediated immune modulation through APC cross-dressing. These results provide mechanistic insight into the rational design of EVs as a cell-free, yet precision medicine-based approach to immunotherapy that can be tailored to promote antigen-specific immune tolerance or pro-inflammatory responses.

## Introduction

Extracellular vesicles (EVs) are small, biologically active vesicles secreted by most cell types that either derive from the endocytic compartment or are shed from the cell membrane (*1*). Small EVs (30-150 nm), which are often referred to as exosomes (*2, 3*), play important roles as messengers carrying information through proteins, nucleic acids, lipids, and metabolites that exert effects on interacting cells (*4-7*). EV membranes contain lipid rafts that are enriched in cholesterol, sphingomyelin, and ceramide, making them highly stable in body fluids (*8*) and thus, an attractive tool for therapeutics. Furthermore, EVs are a source of peptide-MHC (pMHC) and can interact with the immune system in multiple ways in an antigen-specific manner (*9*). Given these advantages, EVs present an attractive tool modular for novel therapeutic development and can be altered in many cases to complement or enhance their potential therapeutic applicability (*10-13*).

Our approach to EV engineering is inspired by the role of endogenous EVs in disease settings such as cancer or autoimmunity (*14, 15*). For example, tumor cell exosome-presented programmed death-ligand 1 (PD-L1) regulates T cell responses in cancer (*16, 17*). Binding of PD-L1 to its receptor PD-1 on T cells results in strong countering of T cell receptor (TCR) signal transduction and CD28 co-stimulation (*18, 19*), thereby suppressing antigen driven activation of T cells. Removal of PD-L1 specifically from exosomes leads to strong T cell activation and tumor rejection (*20*). In autoimmunity, EVs have been shown to potentially contribute to disease progression through immune activation. For example, in type 1 diabetes (T1D), islet autoantigens such as insulin, glutamic acid decarboxylase 65 (GAD65), and insulinoma-associated protein-2 (IA-2) have been identified in beta cell EVs, which can then be transferred to antigen presenting cells (APCs) for processing and presentation to T cells (*21-23*). Many dysregulated immune pathways in cancer and autoimmunity may be modulated, at least in part, by contributions from EVs (*14, 15*). Determining the mechanisms by which EVs impart these effects may provide insights into how to effectively engineer and leverage them in an *ex vivo* setting for therapeutic purposes.

Here, we investigated the negative regulatory properties of engineered EVs as a tool for modulating T cell activation and cytotoxicity to combat the autoimmune component of T1D. We designed an *in vitro* system for generating and testing EVs carrying autoantigen-loaded HLA class I molecules using the T1D risk-associated HLA-A*02, plus co-stimulation with or without PD-L1 co-inhibition. We demonstrated the importance of APCs in EV-mediated activation of CD8^+^ T cells, as well as a potential for CD80 in both EV-mediated stimulation and inhibition. We also generated islet-autoreactive human CD8^+^ T cell “avatars” and showed that immune engineered EVs reduced T cell-mediated killing of target cells *in vitro*. Our results provide valuable insight into effective EV engineering with applications for tissue-specific immune modulation.

## Results

### EV isolation and characterization

When determining a suitable system to use for generating EVs and investigating their immunomodulatory effects, we chose to engineer an *in vitro* system with relevance to T1D. Insulin is one of the primary autoantigens in T1D (*24*), with several pre-proinsulin peptides inducing CD8^+^ T cell mediated autoimmunity through HLA-A*02 presentation (*25*). K562 cells, a lymphoblast cell line widely used as a backbone for artificial antigen presenting cells (aAPCs) (*26*), modified to express CD80, CD83, HLA-A*02 and pre-proinsulin (K562 A2/PPI) were used as the cell source for generating EVs (Supplemental Figure 1). THP-1 cells, which are a monocytic cell line that expresses HLA-A*02 but not CD80 (Supplementary Figure 2A), were also used for generating EVs.

**Figure 1.**
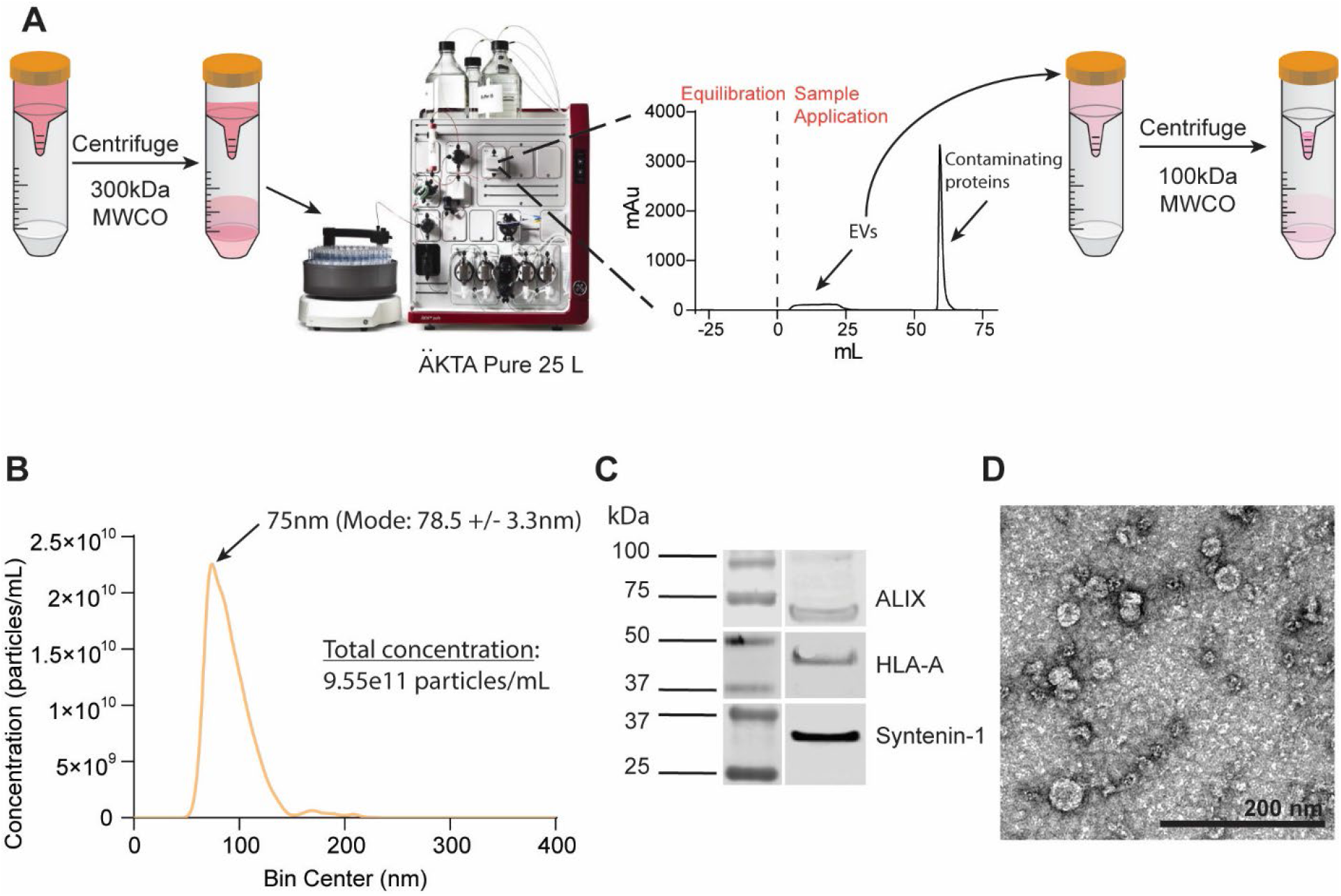
EV isolation and characterization. (A) Schematic overview of the workflow. Conditioned, processed media is filtered through 300kDa MWCO UF devices to remove soluble proteins and concentrate the EV fraction. Concentrated media is loaded onto a BE-SEC column using the ÄKTA Pure chromatography system. The first eluting fractions (EVs) are collected via an automated fraction collector then subsequently concentrated using 100kDa MWCO UF devices and used for further experiments. Contaminating proteins trapped by the BE-SEC column are eluted out during the column wash. (B) NTA showing representative particle concentration and size distribution of EVs derived from K562 A2/PPI cells. (C) Western blot analysis of EV isolates demonstrating the presence of small EV markers. n=3 (D) Representative TEM image of K562 A2/PPI cell-derived EVs. Scale bar = 200nm.

**Figure 2.**
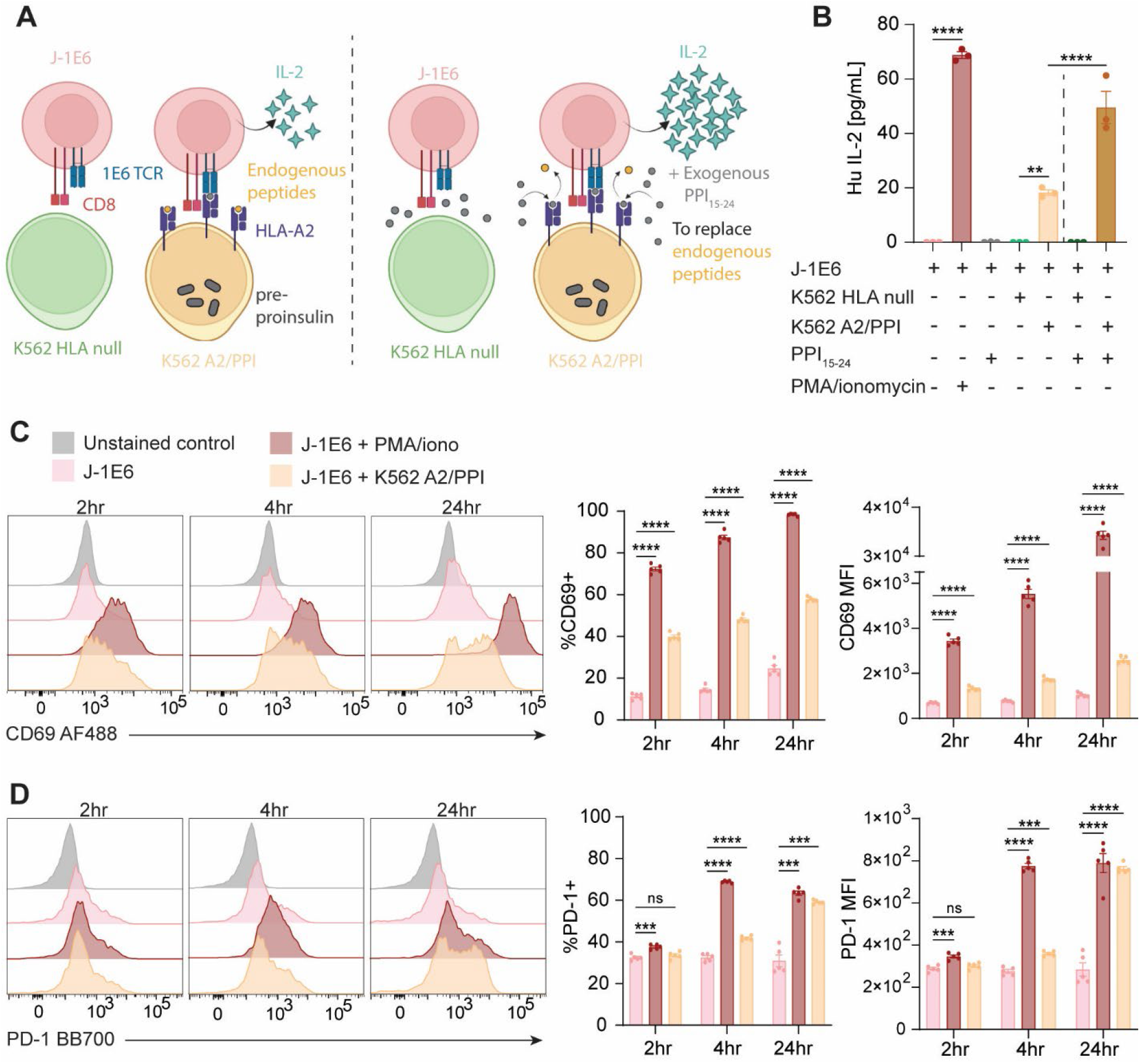
Validating an in vitro target:effector co-culture model for EV engineering. (A) Schematic representation of J-1E6 and K562 co-culture. (B) IL-2 secretion from J-1E6 T cells showing activation from K562 A2/PPI cells, which is increased with exogenous peptide loading in HLA-A*02. K562 cells lacking HLA-A*02 fail to induce IL-2 secretion from J-1E6 T cells even in the presence of exogenous peptide. Representative of n = 4 independent experiments. (C) Surface expression of the activation marker CD69 on J-1E6 T cells over a 24 hour co-culture with K562 A2/PPI cells. n = 5. (D) Surface expression of PD-1 on J-1E6 T cells over a 24 hour co-culture with K562 A2/PPI cells. n = 5. Statistical differences for (B) were determined by one-way ANOVA followed by Tukey’s multiple comparisons test. Statistical differences for (C) and (D) were determined by two-way ANOVA followed by Tukey’s multiple comparisons test. ns = non-significant, ** p < 0.01, *** p < 0.001, **** p < 0.0001. Panel (A) was created using Biorender.com.

While ultracentrifugation has been a common method for small EV isolation (*27*), this can lead to vesicle damage and aggregation (*28, 29*). One alternative method that avoids this potential limitation uses liquid chromatography columns for bind-elute size exclusion separation (BE-SEC) of small EVs from conditioned medium (*30*). We adopted this method, in combination with ultrafiltration steps before and after chromatography separation, to isolate EVs for use in functional assays (Figure 1A). NTA of EVs from both K562 A2/PPI (K-EV) and THP-1 (T-EV) cells isolated through chromatography were consistent with those isolated by ultracentrifugation and consistently showed a homogeneous particle size distribution, with diameters across multiple isolations between 40 and 110 nm (Figure 1B, Supplementary Figure 3A). Western blot analysis demonstrated the presence of small EV markers ALIX and Sytenin-1 (*31*) in both K-EVs and T-EVs (Figure 1C, Supplementary Figure 3B). We were also able to detect HLA-A in EVs from both cell lines (Figure 1C, Supplementary Figure 3B), indicating that HLA is trafficked to EVs from parent cells. TEM imaging of K-EVs showed particles with typical size and cup-shaped morphology (*32*) (Figure 1D). These data together demonstrate that K562 A2/PPI cells and THP-1 monocytes secrete small EVs, which we can purify from conditioned medium in a scalable and reproducible manner.

**Figure 3.**
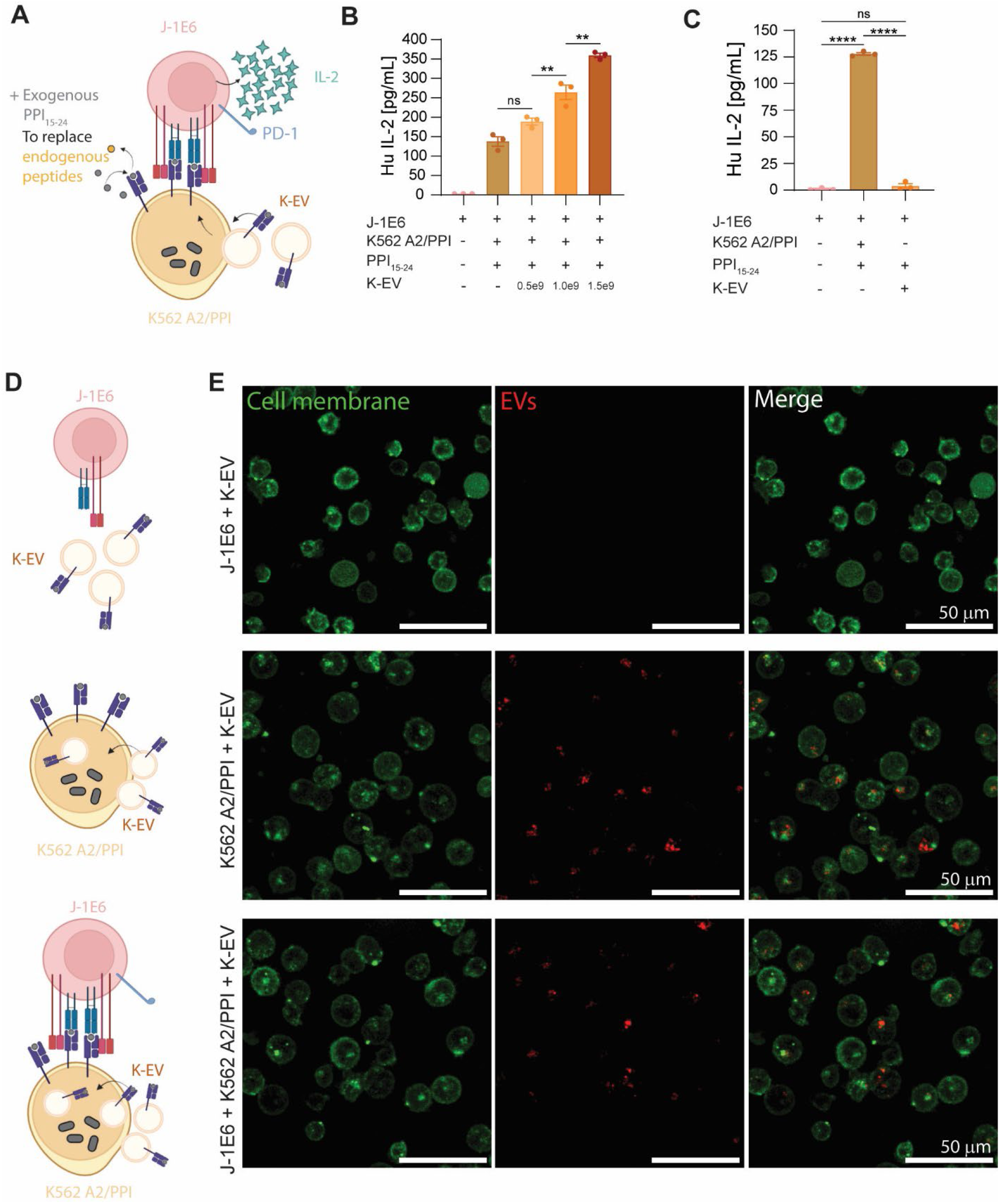
EVs need APCs to modulate T cell activation in vitro. (A) Schematic representation of J-1E6 co-culture with K562 A2/PPI cells and K-EVs. (B) IL-2 secretion from J-1E6 T cells showing increased activation with increasing amounts of K-EVs in cultures. Representative of n = 5 independent experiments. (C) IL-2 secretion from J-1E6 cells after co-culture with K562 A2/PPI cells or EVs, showing that EVs alone do not activate T cells. Representative of n = 4 independent experiments. (D) Schematic representations of the different conditions used in (E) to determine EV interactions in the co-culture system. (E) Confocal microscopy images of DiR-labeled K-EVs incubated with either J-1E6, K562 A2/PPI, or both cell types for 24 hours, after which cell membranes were labeled with MemBrite Fix 568/580. DiR signal was detected in cultures with K562 A2/PPI cells and J-1E6 with K562 A2/PPI cells, but not J-1E6 cells alone, suggesting that K562 A2/PPI cells endocytose EVs. Statistical differences for (B) and (C) were determined by one-way ANOVA followed by Tukey’s multiple comparisons test. ns = non-significant, ** p < 0.01, **** p < 0.0001. Panels (A) and (D) were created using Biorender.com.

### Validating an in vitro target:effector co-culture model for EV engineering

Prior to investigating the immunomodulatory effects of EVs, we first validated that K562 A2/PPI aAPCs, the cells used for manufacturing K-EVs, could activate cognate T cells *in vitro*. Jurkat T cells were engineered as model effector cells to express CD8 and the 1E6 TCR (hereafter referred to as J-1E6 T cells), which recognizes PPI_15-24_ presented in HLA-A*02 (*33*). To validate that K562 A2/PPI cells could activate J-1E6 cells, we performed co-cultures using IL-2 secretion as a readout of T cell antigen recognition and activation (*20*). K562 cells devoid of any modifications (K562 HLA null) were used as negative controls to ensure that T cell IL-2 secretion was dependent on HLA-presented peptide (Figure 2A). T cells did not secrete any IL-2 when co-cultured with K562 HLA null cells even with addition of exogenous PPI_15-24_ peptide, or when cultured with exogenous peptide in the absence of K562 cells (Figure 2B). Conversely, T cells cultured with K562 A2/PPI cells secreted IL-2, albeit to a low extent compared to positive control cells activated with PMA and ionomycin (Figure 2B). Adding exogenous PPI_15-24_ peptide to co-cultures with T cells and K562 A2/PPI cells significantly increased IL-2 secretion above similar cultures without exogenous peptide addition (Figure 2B). K562 A2/PPI cells loaded with a different islet antigen peptide, IGRP_265-273,_ failed to induce IL-2 secretion from T cells above control levels without exogenous PPI loading (Supplementary Figure 4). THP-1 cells, which express HLA-A*02 but not CD80, were cultured with J-1E6 cells in a similar manner with or without exogenous PPI_15-24_ peptide. THP-1 cells failed to induce IL-2 secretion from T cells even with peptide loading, indicating the necessity of CD80 co-stimulation in this model (Supplementary Figure 2B). We also performed flow cytometry to evaluate cell surface markers of activation on J-1E6 cells after 2, 4, and 24 hours of co-culture with K562 A2/PPI cells. J-1E6 T cells co-cultured with K562 A2/PPI cells expressed significantly more CD69 (Figure 2C) and PD-1 (Figure 2D) compared to T cells alone. These results indicate that J-1E6 T cells recognize and react in response to PPI_15-24_ presented in HLA-A*02 on K562 A2/PPI cells and provide a foundation for investigating the immunomodulatory effects of EVs generated from these model cells.

**Figure 4.**
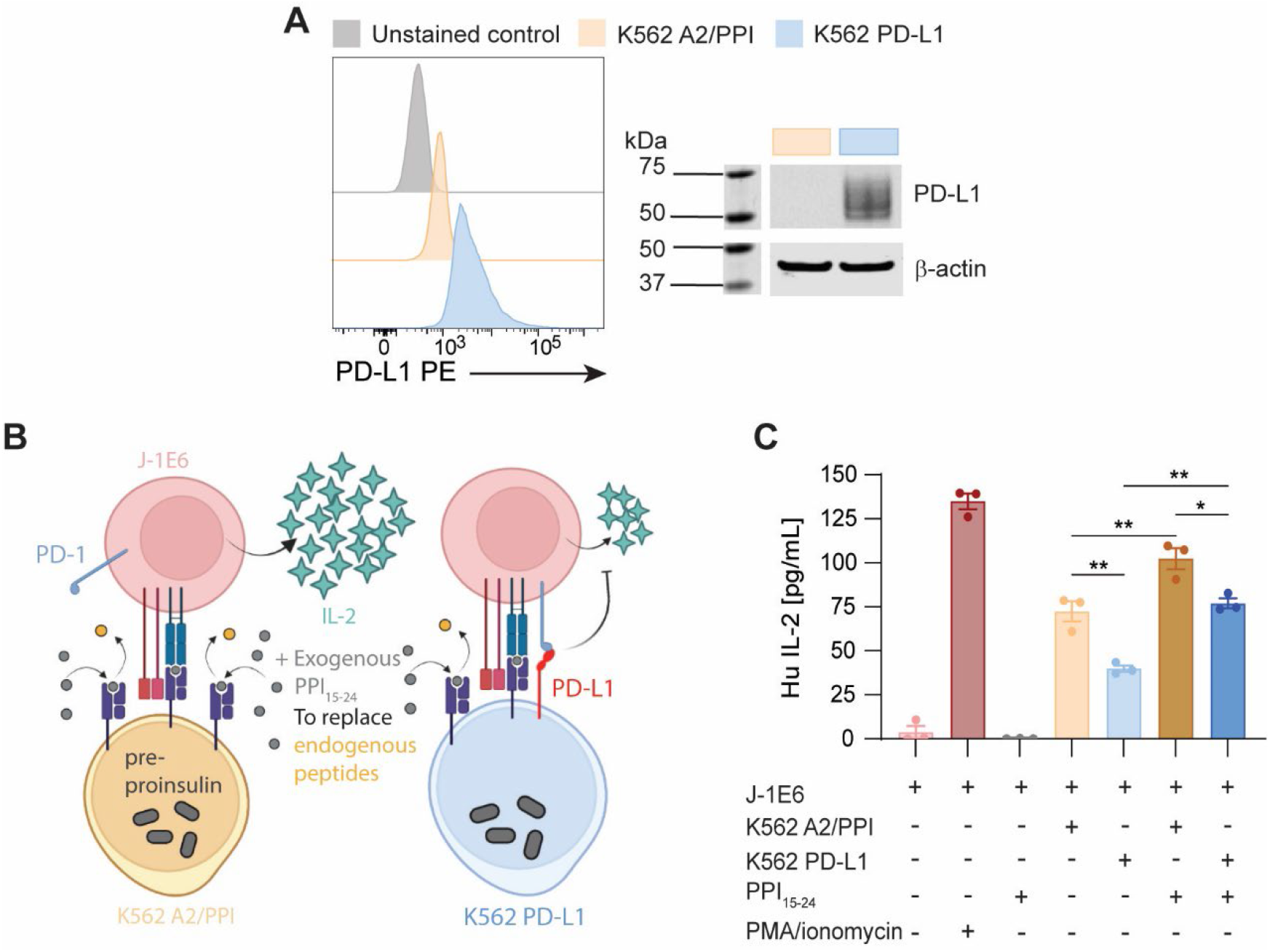
Driving PD-L1 expression in EV parent cells. (A) Surface expression of PD-L1 on lentivirus-transduced cells shown via flow cytometry, and further confirmed by western blot analysis. (B) Schematic representation of J-1E6 co-culture with either K562 A2/PPI or K562 PD-L1 cells, where PD-L1 engages with PD-1 to suppress IL-2 secretion. (C) IL-2 secretion from J-1E6 T cells showing decreased activation when co-cultured with K562 PD-L1 cells compared to K562 A2/PPI. This suppression is still seen when exogenous PPI_15-24_ is added to co-cultures. Representative of n = 9 independent experiments. Statistical differences for (C) were determined by one-way ANOVA followed by Tukey’s multiple comparison test. * p < 0.05, ** p < 0.01. Panel was created using Biorender.com.

### EVs need APCs to modulate T cell activation in vitro

Having established our EV source and *in vitro* model, we next examined the effects of EVs on T cell activation, as there is evidence showing that EVs can influence T cells through a variety of mechanisms including direct T cell contact (*34, 35*) or cross-dressing of APCs (*36, 37*). Cross-dressing refers to the passage of EV pHLA and other molecules to an APC without further antigen processing. We added increasing numbers of K-EVs to co-cultures with T cells and K562 A2/PPI cells (Figure 3A) and measured IL-2 secretion after 24 hours. Addition of K-EVs caused an increase in IL-2 secretion in a dose-dependent manner (Figure 3B). However, the addition of K-EVs to J-1E6 T cells in the absence of aAPCs did not result in significant IL-2 induction when compared to T cells alone (Figure 3C). To further examine the mechanism of EV immune activity, we labeled EVs with DiR and added them to cultures with either J-1E6 T cells alone, K562 A2/PPI cells alone, or J-1E6 and K562 A2/PPI cells together (Figure 3D). After 24 hours, we labeled cell membranes and performed confocal microscopy. Intracellular DiR signal was detectable in K562 A2/PPI cells alone or in co-culture with T cells, but EV uptake was undetectable for J-1E6 T cells alone (Figure 3E). This result suggests that K-EVs pass their contents to K562 A2/PPI cells but do not interact directly with J-1E6 T cells. Adding T-EVs, which contain pHLA but not CD80, to co-cultures of T and K562 A2/PPI cells failed to increase IL-2 secretion (Supplementary Figure 5). This result indicates that EV CD80 is important for EV immune activity in our model despite endogenous CD80 expression in the aAPC. Taken together, these data show that EVs in our system containing pHLA and co-stimulatory molecules from parent cells require aAPCs to impart effects on T cells. Further, these effects depend on co-stimulatory molecules such as CD80 on EV parent cells and thus EVs, as T-EVs containing pHLA but not CD80 fail to activate T cells in the presence of aAPCs.

**Figure 5.**
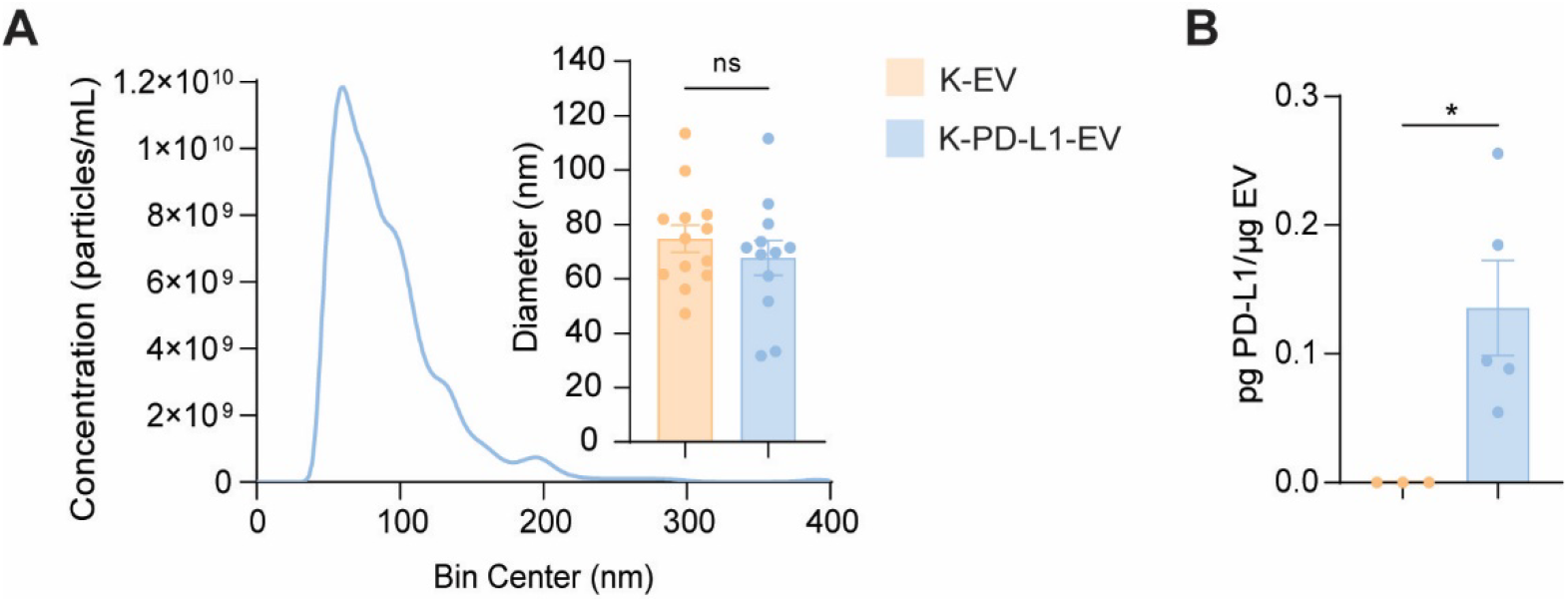
Validating PD-L1 expression in EVs. (A) NTA showing representative particle concentration and size distribution of EVs derived from K562 PD-L1 cells. The inset shows the mode particle diameter for multiple K-EV and K-PD-L1-EV preparations, indicating that lentiviral transduction did not negatively impact EV generation. n = 13 and n = 12. (B) ELISA of PD-L1 on EVs from K562 cells compared to total EV protein content. Data from n = 2 independent experiments. Statistical differences for (A) and (B) were determined by an unpaired, two-tailed t-test. ns = non-significant, * p < 0.05.

### Driving PD-L1 expression in EV parent cells

EV PD-L1 incorporation correlates with cellular expression levels, as cells naturally shuttle the protein to EV membranes (*38*). We therefore reasoned that we could purify PD-L1^+^ EVs from PD-L1 expressing cells. While many cells such as THP-1 monocytes upregulate PD-L1 expression in the presence of inflammatory cytokines like IFN-γ (*39-41*) (Supplementary Figure 6A), we chose to create a K562 A2/PPI cell line stably expressing PD-L1 through lentiviral transduction. After lentiviral transduction, we sorted and expanded PD-L1^+^ K562 A2/PPI cells (K562 PD-L1), then confirmed stable PD-L1 expression via flow cytometry and western blot after several passages (Figure 4A). Recent work has highlighted the influence of *cis*-PD-L1/CD80 binding and its inhibitory effect on these molecules binding in *trans* to PD-1 and CD28 or CTLA-4 (*42, 43*). While CD80 typically exists as a homodimer in cell membranes, its affinity for PD-L1 is higher than it is for itself (*44*), which can lead to preferential formation of PD-L1/CD80 heterodimers (i.e., *cis*-PD-L1/CD80). These heterodimers block PD-L1 from binding to PD-1 on other cell membranes, thus inhibiting *trans*-PD-L1/PD-1 signaling (*43*). In particular, high CD80 expression can lead to increased *cis*-PD-L1/CD80 heterodimers and subsequent decreased *trans*-PD-L1/PD-1 binding. As such, our next step was to ensure that K562 PD-L1 cells would suppress T cell activation in our co-culture model (Figure 4B). We cultured J-1E6 T cells with K562 A2/PPI or K562 PD-L1 cells, with or without exogenous PPI_15-24_ peptide for 24 hours and measured IL-2 secretion in the supernatant. Co-culture of T cells and K562 PD-L1 cells resulted in less IL-2 production as compared to co-culture with K562 A2/PPI cells, even with exogenous peptide (Figure 4C). The results shown here indicate K562 cells presenting PD-L1 alongside cognate pHLA complexes can reduce T cell activation in our model, despite high CD80 expression on K562 aAPCs.

**Figure 6.**
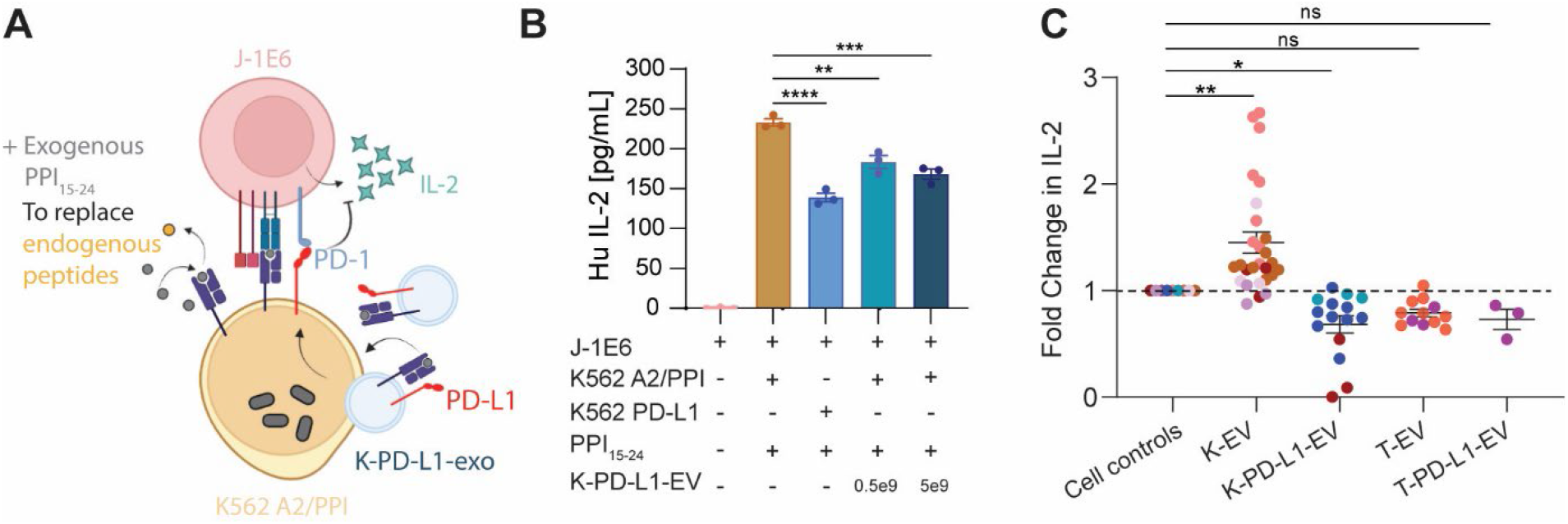
K-PD-L1-EVs suppress T cell activation. (A) Schematic representation of J-1E6 co-cultures with K562 A2/PPI cells and K-PD-L1-EVs. (B) IL-2 secretion from J-1E6 cells showing decreased activation with K562 PD-L1 cells or increasing amounts of K-PD-L1-EVs in cultures. Fold change in IL-2 secretion across all experiments with different exosome treatments, showing that K-EVs significantly increase and K-PD-L1-EVs significantly decrease IL-2 secretion. T-EVs and T-PD-L1-EVs have no significant effect. Each dot represents a technical replicate from an independent experiment, and each color represents an independent experiment. Each technical replicate was normalized to its independent experimental control with just J-1E6 and K562 A2/PPI cells. Statistical differences for (B) were determined by one-way ANOVA followed by Tukey’s multiple comparisons test. Statistical differences for (C) were determined using a linear mixed effects model where fold change in IL-2 is predicted by the EV treatment as a linear fixed effect and the individual experiment as a random effect. ns = non-significant, * p < 0.05, ** p < 0.01, *** p < 0.001, **** p < 0.0001. Panel (A) was created using Biorender.com.

### Validating PD-L1 expression in EVs

We next sought to characterize EVs from K562 PD-L1 cells (K-PD-L1-EV). NTA analysis of K-PD-L1-EVs showed a homogeneous size distribution with mean particle diameters similar to K-EVs (Figure 5A). Further, we detected PD-L1 in EV isolates through an ELISA (Figure 5B). We also detected PD-L1 in EVs from THP-1 cells cultured with IFN-γ (T-PD-L1-EV, Supplementary Figure 6B). These results demonstrate that stable expression of PD-L1 in parent cells naturally leads to packaging of PD-L1 into EVs, which is detectable together with HLA.

### K-PD-L1-EVs suppress T cell activation

The expression of PD-L1 on EVs in circulation is associated with poor outcomes of anti-PD-1 cancer therapies (*17*) due to their ability to suppress cytotoxic T cells that would normally eradicate tumor cells. Removal of PD-L1 specifically in EVs restores anti-tumor immunity (*20*). To examine if reciprocal effects can be observed in the context of the autoimmune disease T1D, we tested EVs with PD-L1 in our *in vitro* target:effector co-culture model (Figure 6A). Increasing amounts of K-PD-L1-EVs in co-culture with T cells and K562 A2/PPI cells led to a decrease in IL-2 secretion in a dose-dependent manner (Figure 6B). Given that these co-cultures were performed with aAPCs completely devoid of PD-L1, these results show that EV PD-L1 is capable of inhibiting antigen recognition-induced activation of T cells specific for a T1D autoantigen. However, this seems to depend on the presence of CD80 on the EVs, as adding T-PD-L1-EVs without CD80 to co-cultures resulted in only a modest decrease in IL-2 secretion from J-1E6 cells that was similar in magnitude to cultures with a large dose of T-EVs (Supplementary Figure 6C).

We performed an analysis across all experiments with EV treatments and compared the fold change in IL-2 secretion (Figure 6C). EVs from K562 cells with or without PD-L1 significantly decreased and increased IL-2 secretion, respectively. EVs from THP-1 cells did not have a significant effect on IL-2 secretion. This analysis provides further evidence that regardless of HLA or PD-L1 expression in EVs, the cell source and additional co-stimulatory markers are important to impart immunomodulatory effects.

### EV PD-L1 reduces T cell-mediated target cell killing in vitro

While Jurkat T cells are a suitable cell line to use for proof-of-principle studies with therapies intended to modulate T cell functionality, they are not true effector cells and thus, lack any appreciable cytotoxic capabilities. We therefore replaced J-1E6 cells with a more authentic effector T cell in our co-culture model (Figure 7A) to determine if PD-L1 presenting EVs still exert their suppressive effects, while keeping our studies within the context of T1D. Primary human CD8^+^ T cell “avatars” were generated to express a TCR specific for the IGRP peptide IGRP_265-273_ presented in HLA-A*02, which is a known autoantigen in the human T1D population (*45*) (referred to here as T-IGRP cells). K562 aAPCs (either K562 A2/PPI or K562 PD-L1) were loaded with exogenous IGRP_265-273_ peptide prior to co-culture with T-IGRP cells and/or K-PD-L1-EVs to provide T cell recognition of cognate pHLA complex. T cell-mediated killing of IGRP loaded K562 target cells was determined after 16 hours by flow cytometry with AV and PI staining. There was minimal K562 cell death in the absence of any T cells. IGRP_265-273_ loaded K562 A2/PPI cells cultured with T-IGRP cells had significantly more AV^+^/PI^+^ cells than those cultured without T-IGRP cells. K562 PD-L1 cells were slightly resistant to T cell-mediated killing, with a 20% reduction in AV^+^/PI^+^ cells compared to K562 A2/PPI cells (Figure 7B). When K562 A2/PPI cells were co-cultured with T-IGRP cells and PD-L1 EVs, the percentage of AV^+^/PI^+^ cells was further reduced to approximately 40% of that in cultures without EVs (Figure 7B). This indicates that EV PD-L1 inhibited antigen-specific T cell-mediated killing of target cells, independent of any cell surface-presented PD-L1. In sum, these data suggest that EVs engineered to present pHLA and PD-L1 can promote immune tolerance against beta cells for T1D immunotherapy.

**Figure 7.**
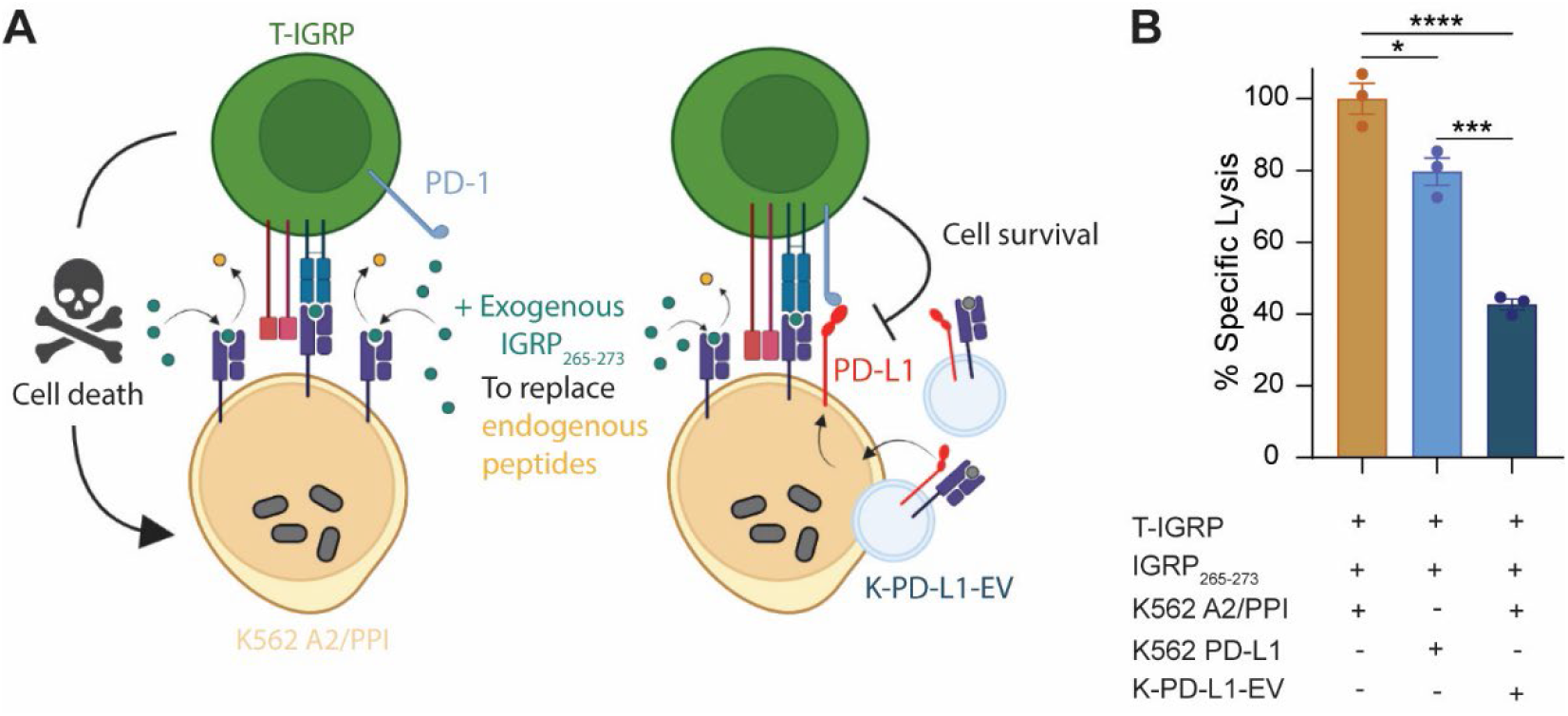
EV PD-L1 reduces T cell-mediated target cell killing in vitro. (A) Schematic representation of T-IGRP co-cultures with K562 A2/PPI cells and K-PD-L1-EVs. (B) Specific lysis of K562 cells by T-IGRP avatars, showing a 60% reduction in specific lysis of K562 A2/PPI cells when K-PD-L1-EVs are added to co-cultures. Statistical differences for (B) were determined by one-way ANOVA followed by Tukey’s multiple comparisons test. * p < 0.05, *** p < 0.001, **** p < 0.0001. Panel (A) was created using Biorender.com.

## Discussion

EVs represent a potential therapeutic option for autoimmune diseases to impart many of the advantages of adoptively transferred cells while reducing some key limitations including cost, scalability, and patient risks from the transfer of living cells (*46-48*). While there is abundant evidence that EVs influence immune outcomes (*9*), the mechanisms by which they influence T cell fate both *in vitro* and *in vivo* remain subject to debate (*15*). Some groups have shown that direct interactions occur between T cells and EVs *in vitro* (*34, 35*). Guo and colleagues performed a series of studies demonstrating that exosomes can directly suppress T cells through PD-1/PD-L1 interactions, but this was dependent on exosomal ICAM-1 (*49*). Conversely, other studies have shown that at least *in vitro*, APCs are required to stimulate T cells through EV uptake and cross-dressing of EV proteins onto the APC cell surface (*36, 37*), which is a process that has been previously studied (*50, 51*) and supports the mechanism we propose here.

Our study investigated the effects of rationally engineered EVs in modulating T cell activation *in vitro*. We designed and validated a target:effector co-culture system with relevance to T1D, using engineered K562 cells as aAPCs and either modified Jurkat or primary human CD8^+^ T cells as effectors. We showed that Jurkat T cells expressing CD8 and the 1E6 TCR clone respond specifically to PPI_15-24_ presented in HLA-A*02 in the presence of co-stimulation, as THP-1 cells loaded with PPI_15-24_ but lacking CD80 failed to induce IL-2 secretion from these cells. We also engineered PD-L1 into K562 cells and showed that this decreased T cell activation, even in the presence of excess exogenous peptide. Both K562 and THP-1 cells secreted EVs, which we were able to purify and use for functional studies in co-cultures with K562 and Jurkat T cells. In our system, K562 aAPCs internalized EVs during co-culture, and APCs were required for T cell activation by EVs. Together, these data point toward the cross-dressing mechanism, wherein EVs are endocytosed and their cargo is redistributed to the cell membrane for presentation to T cells. Additionally, we provide evidence that EV PD-L1 can inhibit T cell activation through aAPC cross-dressing in the presence of co-delivered CD80.

An interesting observation that arose from our studies was the inability of EVs derived from THP-1 cells to impart the same stimulatory effects as K562-derived EVs in co-cultures with Jurkat T cells and K562 aAPCs, despite the presence of HLA on both EV populations, and excess exogenous PPI_15-24_ in the system. Given the evidence that EVs cross-dressed aAPCs, we suggest that this difference stems from the lack of CD80 on THP-1 cells and T-EVs. A very large dose of T-EVs did result in a modest decrease in IL-2 secretion from J-1E6 cells. We speculate this was due to extensive HLA cross-dressing on K562 aAPCs, thus providing an overabundance of pHLA to T cells in the absence of proportional of CD80 co-stimulation. Thus aAPCs treated with K-EVs are cross-dressed with proportionate amounts of signals 1 (pHLA) and 2 (CD80) needed for T cell activation.

When designing and testing EVs containing PD-L1, we were initially focused on PD-1/PD-L1 interactions and the necessity of pHLA/TCR interactions for PD-1 signaling to occur (*52, 53*). However, the data we have with PD-L1 EVs from K562 and THP-1 cells suggests that CD80 also plays a role in EV-mediated T cell suppression through the APC cross-dressing pathway. The importance of EV CD80 has been demonstrated before, with increased CD80 leading to increased immune activation via dendritic cell cross-dressing (*54*) and decreased CD80 leading to a more regulatory immune state (*55*). Several groups have also shown that PD-L1 and CD80 interact on the same cell membrane to form *cis*-heterodimers, and this influences if and how these molecules can then bind to PD-1 and CD28 or CTLA-4 on other cell membranes (*42, 43*). Okazaki and colleagues have even demonstrated that disrupting *cis*-CD80/PD-L1 interactions alleviates multiple autoimmune disease models (*56*). As Jurkat T cells do not express CTLA-4 (*57, 58*), the interactions taking place in our model system would be between *cis*-CD80/PD-L1 and *trans*-PD-1/PD-L1. EV delivery of HLA to aAPCs remained consistent across all EV conditions with differences lying in co-delivery of PD-L1 and CD80, and only EVs co-delivering HLA, PD-L1, and CD80 able to significantly suppress T cell activation.

Delivery of nanoparticulate pMHC complexes has been previously demonstrated to impact T1D development. Santamaria and colleagues showed that synthetic nanoparticles coated with beta cell pMHC complexes are able to delay or prevent T1D progression in NOD mice (*59, 60*). While these studies demonstrated direct interactions between 40-60 nm nanoparticles and autoreactive T cells (*61*), others have shown that pMHC and anti-CD28 coated nanoparticles under 300 nm are unable to initiate TCR signaling events through such direct interactions (*62*), similar to our observations with small EVs.

Importantly, we showed that PD-L1-containing EVs were able to suppress T cell avatar-mediated, antigen-specific killing of target cells. Future studies will be needed to determine whether these EVs are able to suppress killing of primary beta cells, including whether engineered EVs can be used as therapeutics to impact the progression or initiation of T1D. Of relevance to our study, Eizirik and colleagues have shown that PD-L1 is expressed in insulin containing islets of people during early stages of T1D (*63*), and they as well as Fife and colleagues have demonstrated that inflammatory cytokines increase beta cell PD-L1 expression (*41, 63*). Thus, it may be possible that beta cells naturally release EVs containing autoantigens and PD-L1 during early stages of T1D, which could be an alternative strategy for generating tolerogenic EVs for use in a T1D setting.

The results presented here advance the understanding of the necessary conditions for EV-mediated T cell stimulation or inhibition, as well as considerations for engineering EVs for tissue-specific immune modulation. This approach for immune modulation combines the advantages of facile production of native HLA with the ability to engineer immune-regulatory functions in a plug-and-play fashion and is easily adaptable to other autoimmune diseases through delivery of different autoantigen pHLA complexes in EVs. In summary, EVs are a potentially unlimited source of nanoparticulate, multimeric, personalized HLA that can be both mass-produced and further engineered to fine-tune immunomodulatory capacity.

## Methods

### Cell culture

The chronic human mylogenous leukemia K562 cell line (*64*) (HLA null) was obtained from ATCC. K562 cells expressing HLA-A2, CD80, and CD83 were a gift from Drs. James Riley and Bruce Levine (University of Pennsylvania) and transduced with a lentivirus construct to express pre-proinsulin (PPI) in order to generate the K562 A2/PPI line. K562 A2/PPI cells were further transduced with a lentivirus construct to express PD-L1 in order to generate K562 PD-L1 cells. Jurkat cells (ATCC, clone E6-1; TIB152™), a human T cell line (*65*), were transduced with lentiviral constructs containing CD8 and the PPI_15-24_-reactive TCR (clone 1E6) (*33*). K562 and Jurkat cell lines were cultured in complete RPMI medium (cRPMI; RPMI-1640 containing L-glutamine supplemented with 10% fetal bovine serum [FBS], 1% penicillin-streptomycin, 10 mM HEPES, 1 mM sodium pyruvate, 1X non-essential amino acids, 0.2 μM 2-mercaptoethanol, final pH 7.4) and maintained at 37°C, 5% CO_2_ atmosphere. THP-1 cells, a human acute monocytic leukemia cell line (*66*), were obtained from ATCC and maintained in RPMI containing L-glutamine supplemented with 10% FBS, 1% penicillin-streptomycin, 10 mM HEPES, 1 mM sodium pyruvate, 50 μM 2-mercaptoethanol, and 4.5 g/l glucose. THP-1 cells were supplemented with 100 U/ml recombinant human IFN-γ (BioLegend) for 3 days to induce PD-L1 expression. To generate EVs, K562 or THP-1 cells were cultured in T-150 flasks in their respective mediums, described above, made with EV-depleted FBS, which was prepared by centrifuging the FBS for 16 hours at 100,000 x g in a Beckman Coulter Optima L-90K Ultracentrifuge at 4°C. Medium was collected and replaced every 3 days, and cells were kept below 80% confluency. Conditioned medium from cells was stored at -80°C until EV isolation.

### EV isolation via chromatography

Conditioned medium was thawed and centrifuged at 300 x g for 5 minutes to pellet cell debris. Supernatant was transferred to fresh tubes and centrifuged at 2,000 x g for 10 minutes to pellet larger extracellular particles. The supernatant was then passed through a 0.22 μm vacuum filter. Large volumes (up to 300 ml) of conditioned medium were concentrated through Sartorius Vivaspin 300 kDa molecular weight cut-off (MWCO) spin-filters at 2,500 x g for 3 minutes to a volume of less than 20 ml. Filtrate was discarded, and retentate was removed and saved in a separate tube. The concentrated conditioned medium was then loaded onto a HiScreen Capto Core 700 column connected to an ÄKTA Pure 25 L Chromatography system (Cytiva Life Sciences). Flow rate settings for column equilibration, sample loading, and column clean in place procedure were chosen according to the manufacturer’s recommendations. The EV fraction was collected according to the 280 nm UV absorbance chromatogram and concentrated using Amicon Ultra-15 100 kDa MWCO spin-filters. Concentrated fractions were washed with 15 ml PBS in the same spin-filter and concentrated to a final volume of 150 μl. All spins and chromatography were performed at 4°C. Samples were stored at 4°C for no more than one week prior to use in functional assays.

### EV isolation via ultracentrifugation

Conditioned medium was thawed and centrifuged at 300 x g for 5 minutes to pellet cell debris. Supernatant was transferred to fresh tubes and centrifuged at 2,000 x g for 10 minutes and at 16,500 x g for 20 minutes, followed by filtration through a 0.22 μm vacuum filter. EVs were pelleted by ultracentrifugation at 110,000 x g for 70 minutes in a Beckman Coulter Optima L-90K Ultracentrifuge. Pellets were resuspended in 1 mL PBS to wash and pelleted again at 120,000 x g for 75 minutes. After the wash, EVs pellets were resuspended in 150 μL PBS and stored until use. All spins were performed at 4°C. For short-term storage (< 1 week), samples were stored at 4°C. For longer-term storage, samples were stored at - 80°C. All EVs isolated through ultracentrifugation were used for analysis of protein content and not functional studies.

### Nanoparticle tracking analysis

Nanoparticle tracking analysis (NTA) was performed using a Nanosight LM14 (Malvern Instruments) consistent with methods previously described(*67*). EVs were diluted in PBS to a final concentration between 1:250 and 1:1000 prior to measurement to fit into the resolution window recommended by the manufacturer. Samples were loaded into the sample stage and the camera level was adjusted to 11 or 12. Three captures of 60 seconds each were recorded for each sample. Nanosight NTA 3.1 software was used to analyze particle size and concentration with a particle detection threshold at 2 or 3. Average particle concentrations measured by the software were multiplied by the dilution factor to obtain the final particle concentration for each sample.

### Electron microscopy

EVs were examined by transmission electron microscopy negative stain at the UF Interdisciplinary Center for Biotechnology Research (ICBR) Electron Microscopy Core, RRID: SCR_019146. Poly-L-lysine treated, 400 mesh carbon coated Formvar copper grid was floated onto 3 μl of aliquoted EV suspension for 5 minutes, incubated on 2% paraformaldehyde in PBS (pH: 7.20) for 15 minutes, followed by a PBS wash and water wash for 5 minutes, each. Excess solution was drawn off with filter paper, and the grid was floated onto 1% aqueous uranyl acetate for 30 seconds. The stain was removed with filter paper, air dried and examined with a FEI Tecnai G2 Spirit Twin TEM (FEI Corp., Hillsboro, OR) operated at 120 kV. Digital images were acquired with a Gatan UltraScan 2k x 2k camera and Digital Micrograph software (Gatan Inc., Pleasanton, CA).

### Western Blot Analysis

Total protein was extracted from cells using RIPA lysis buffer supplemented with Pierce Protease inhibitor tablets (Thermo Fisher Scientific). Protein content in cell lysates and EVs was determined using the Pierce BCA Protein Assay Kit (Thermo Fisher Scientific). Between 5 and 20 μg of total protein (kept consistent for individual experiments) was loaded into a 4-12% Bis-Tris gel, resolved by SDS-PAGE under reducing conditions, and transferred to a PVDF membrane. Membranes were blocked with Intercept PBS Blocking Buffer (Licor) for 1 hour, followed by incubation with primary antibodies overnight at 4°C. Membranes were washed 4 times for 5 minutes each with PBS + 0.1% Tween 20, incubated with IRDye^©^ secondary antibodies (Licor) for 1 hour at room temperature and washed an additional 4 times. All blocking and antibody incubations were performed with gentle rocking. Membranes were imaged using a Licor Odyssey Infrared Western Block Scanner. Antibody information is listed in Supplementary Table 1.

### Lentiviral transduction of K562 and Jurkat cells

K562 cells expressing HLA-A2, CD80, and CD83 were transduced with lentiviral vectors to express human PPI and PD-L1. Pre-packaged PD-L1 lentivirus containing puromycin resistance was purchased from Vigene Biosciences (LH872950). PPI lentivirus was generated in-house from human PPI plasmid, which was a gift from Andrew Sewell (Cardiff University) and Mark Peakman (King’s College London). Lentiviral vectors were used to express CD8 and the PPI-1E6 TCR (*68*) (both generated in-house) in human Jurkat T cells (J-1E6). Generation of all lentivirus constructs was performed as previously described (*69*). Transductions were carried out according to previously published protocols (*69*). Briefly, K562 or Jurkat T cells were seeded into a 24-well plate at 2.5 × 10^5^ cells/well in 1 ml cRPMI and transduced in the presence of protamine sulfate (8 μg/ml, Sigma-Aldrich). Transgene expression was assessed 72 hours post-transduction by flow cytometry. To generate a stable PD-L1 expressing K562 cell line (K562 PD-L1), PD-L1^+^ cells were sorted for on a BD FACS Aria III by PD-L1 surface staining (PD-L1 PE, clone 29E.2A3, BioLegend), then expanded *in vitro* in T-150 flasks in cRPMI for another week in the presence of 2 μg/ml puromycin before any functional studies or EV isolations.

### Flow cytometry

Cells were stained with Live/Dead Near-IR Fixable Viability Dye (Thermo Fisher) according to the manufacturer’s instructions. Next, cells were washed with FACS buffer (PBS + 2% FBS + 0.05% NaN_3_) and incubated with Fc receptor blocking solution (Human TruStain FcX, BioLegend) for 5 minutes. Fluorophore conjugated antibodies (Supplementary Table 1) were directly added and incubated for 30 minutes in the dark at 4°C. Cells were washed twice and resuspended in FACS buffer and passed through a 35 μm strainer just before acquisition on a BD FACSCelesta flow cytometer. Marker positivity was set using single color-stained UltraComp eBeads (Thermo Fisher Scientific) and fluorescence minus one stained control cells. Analyses were performed using FlowJo software v10.6.2 (BD Life Sciences).

### Extracellular vesicle PD-L1 analysis

PD-L1 was detected in K562 or THP-1 EVs using the Human PD-L1 DuoSet ELISA kit and DuoSet ELISA Ancillary Reagent Kit 2 (R&D Systems) according to the manufacturer’s instructions. EVs were lysed prior to analysis using 1% Triton-X 100, and total protein content was measured using the Pierce BCA Protein Assay Kit.

### Jurkat stimulation

K562 cells (either HLA null, A2/PPI, or PD-L1) were seeded in U-bottom 96 well plates at 1 × 10^5^ cells/well in 100 μl cRPMI, then cultured for 4 hours in the presence of PPI_15-24_ (ALWGPDPAAA; kindly provided by Holger Russ, University of Colorado Denver) or IGRP_265-273_ (VLFGLGFAI; Genscript) at 10 μg/ml to pre-load with exogenous peptide. Following this, J-1E6 T cells were added at 2 × 10^5^ cells/well, along with any EV treatments (amounts indicated in results and figures), in 100 μl for a final well volume of 200 μl. J-1E6 T cells cultured without K562 and peptide were used as a negative control, and those treated with 5 ng/ml phorbol 12-myristate 13-acetate (PMA) and 0.2 μM ionomycin were used a positive control. In some experiments, Jurkat 1E6 T cells were cultured with 10 μg/ml PPI_15-24_ peptide in the absence of K562 cells as an additional control. Co-cultures were performed in triplicate and incubated for 24 hours at 37°C, 5% CO_2_ atmosphere, after which the culture supernatant was collected and stored at -80°C until analysis. The co-culture experiments were performed similarly with THP-1 cells replacing K562 cells, as indicated in the results section. Human IL-2 in the supernatant was measured using a sandwich ELISA kit (Human IL-2 Quantikine ELISA, R&D Systems) according to the manufacturer’s instructions. The values for fold change in IL-2 secretion were calculated using the following formula:

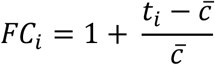

where *FC*_*i*_ is the calculated fold change value, *t*_*i*_ is the measured IL-2 secretion of a technical replicate for a given EV treatment, and 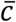is the average IL-2 secretion from the appropriate experimental positive control group.

### Extracellular vesicle uptake assay

EVs from 200 ml of K562 A2/PPI conditioned medium were labeled with 1 μM DiR lipophilic dye (Thermo Fisher Scientific) for 15 minutes at room temperature after the first ultrafiltration step, but immediately prior to chromatography separation. Chromatography and subsequent ultrafiltration were then carried out as described above. K562 cells were seeded into a 12 well-chambered coverslip (ibidi) at 1 × 10^5^ cells/well in 100 μl cRPMI and pre-loaded for 4 hours with 10 μg/ml PPI_15-24_. Following peptide loading, 3 × 10^10^ DiR-labeled EVs with or without 2 × 10^5^ J-1E6 T cells were added to wells containing peptide-loaded K562 cells in 150 μl for a final well volume of 250 μl. As a negative control, 2 × 10^5^ J-1E6 cells and DiR-labeled EVs in 250 μl cRPMI were seeded in wells lacking K562 cells. Cultures were performed in duplicate and incubated for 24 hours at 37°C, 5% CO_2_ atmosphere, after which cell membranes were stained with Membrite Fix 568/580 (Biotium) according to the manufacturer’s instructions. Cells were imaged in the same 12 well-chambered coverslip using a Leica SP8 confocal microscope with HyD detectors, with a 20x/0.75 numerical aperture Plan-Apochromatic air objective.

### Lentivirus production and transduction of primary human T cells

Peripheral blood mononuclear cells (PBMCs) were isolated from leukopheresis-enriched blood from healthy donors (median age: 27 years, range 20-27 years, N=3, 66% female) purchased from LifeSouth Community Blood Center (Gainesville, FL). Buffy coats were processed to isolate PBMCs via Ficoll density gradient centrifugation followed by naïve CD8^+^ T cell isolation with EasySep Human Naïve CD8^+^ T Cell Isolation Kit II (STEMCELL). Lentiviral vector encoding the Islet-specific Glucose-6-Phosphatase Catalytic Subunit 2 (IGRP)-reactive TCR (clone 32, IGRP_265-273_-reactive) was produced as previously described (*70, 71*). Autoreactive CD8^+^ T cell “avatars” expressing this TCR were generated as previously described (*69, 71*). Briefly, isolated naïve CD8^+^ T cells were plated at 2.5 × 10^5^ cells/well in a 24-well plate in 2 ml cRPMI on day 0 and activated with DynaBeads Human T-Activator anti-CD3/CD28 coated microbeads (Thermo Fisher Scientific) and recombinant human IL-2 (rIL-2; NCI Biological Resources Branch Preclinical Repository). After 48 hours incubation at 37C, 5% CO_2_, cells were transduced with 3 transduction units (TU)/cell of lentivirus in the presence of 8 μg/ml protamine sulfate and 100IU rIL-2 and spinnoculated at 1,000 x g for 30 minutes at 32°C. Transduction efficiency was assessed by eGFP expression via flow cytometry 3-5 days post-transduction.

### Cell mediated lysis assay

K562 A2/PPI cells were labeled with CellTrace Violet (CTV, Invitrogen) and seeded in a 48 well plate at 30,000 cells/well in 300 μl cRPMI. Twenty-four hours later, the cells were pre-loaded with 10 μg/ml IGRP_265-273_ peptide for 4 hours. Then, IGRP T cell avatars were added to wells at a 5:1 target:effector ratio, with or without EVs from 300 mL of K562 PD-L1 conditioned medium for a final well volume of 600 μl. Co-cultures were incubated for 16 hours at 37°C, 5% CO_2_, after which cells were collected into FACS tubes and stained with propidium iodide (PI, Invitrogen) and AlexaFluor 647-labeled Annexin-V (AV, BioLegend) in Annexin V binding buffer (BioLegend) according to the manufacturer’s instructions. Cells were acquired on a BD FACSCelesta and analyzed using FlowJo software v10.6.2. The percent specific lysis of CTV^+^ K562 cells was calculated as previously described (*72*).

### Data and statistical analysis

All measurements were taken from distinct biological or technical replicates. Means among three or more groups were compared by one- or two-way analysis of variance (ANOVA) with Tukey’s post-hoc pairwise comparison, and means between two groups were compared by two-tailed Student’s *t* test in GraphPad Prism version 9 software. A linear mixed effects model in RStudio was used to determine statistical significance when comparing fold change in IL-2 across all experiments with EVs. This model was chosen over ANOVA due to the clustered data and unbalanced design across individual experiments. A confidence level of 95% was considered significant. The statistical test used, exact *P* values, and definition of *n* are indicated in the figure legends. Error bars display the mean ± standard error of the mean (SEM).

## Acknowledgements

This work was funded by National Institutes of Health grants F31DK126397 (M.W.B.), T32DK108736 (M.W.B. and L.D.P.), F31DK129004 (L.D.P.), R01DK132387 (E.A.P.), P01AI042288 (T.M.B., E.A.P.), and UH3DK122638 (T.M.B., E.A.P.). Thanks to the UF ICBR Electron Microscopy, RRID:SCR_019146, and Rudolfo Alvarado for performing electron microscopy.

## Author contributions

M.W.B. performed experiments and data analysis. T.M. and L.D.P. provided reagents and discussions critical to the manuscript. T.M. and L.D.P. generated T cell avatars. T.M.B. provided critical expertise and support. M.W.B. and E.A.P. designed the study and wrote the manuscript. All authors discussed the results and commented on the manuscript.

E.A.P. is the guarantor of this work and, as such, had full access to all data in the study and takes responsibility for the integrity of the data and the accuracy of the data analysis.

