## Supplementary Information for "Immune engineered extracellular vesicles to modulate T cell activation in the context of type 1 diabetes"

**Supplementary Table 1.** Antibody information.

| Antigen | Dilution | Host | Supplier | Catalog number | Clone |
|---|---|---|---|---|---|
| CD80 | 1:50 | Mouse | BD | 566263 | L307.4 |
| HLA-A2 | 1:50 | Mouse | BD | 561341 | BB7.2 |
| CD83 | 1:50 | Mouse | BD | 551073 | HB15e |
| PD-L1 | 1:50 | Mouse | BioLegend | 329706 | 29E.2A3 |
| CD8 | 1:50 | Mouse | BioLegend | 301046 | RPA-T8 |
| CD69 | 1:50 | Mouse | BioLegend | 310916 | FN50 |
| PD-1 | 1:50 | Mouse | BD | 566460 | EH12.1 |
| PD-L1 | 1:200 | Rabbit | Cell Signaling | 13694S | E1L3N |
| ALIX | 1:500 | Mouse | Abcam | ab117600 | 3A9 |
| HLA-A | 1:1000 | Rabbit | Abcam | ab52922 | EP1395Y |
| Syntenin-1 | 1:500 | Rabbit | Abcam | ab133267 | EPR8102 |
| beta-actin | 1:10000 | Mouse | Sigma | A1978 | AC-15 |

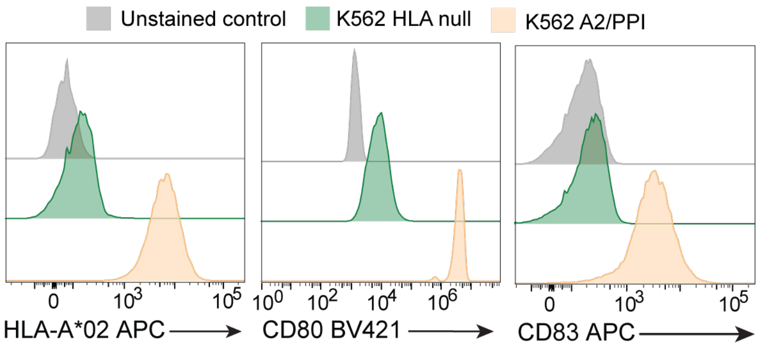


**Supplementary Figure 1.** Surface expression of HLA-A*02, CD80, and CD83 on K562 cell lines, showing strong expression on K562 A2/PPI cells but not K562 HLA null cells. n=3.

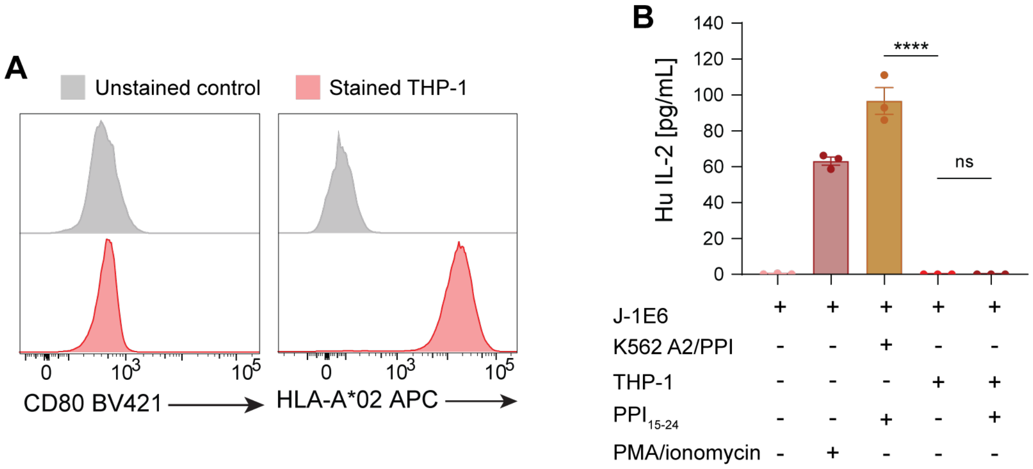


**Supplementary Figure 2**. Characterization of THP-1 cells and their effects in co-cultures with J-1E6 T cells. (A) Surface expression of CD80 and HLA-A*02 in THP-1 cells. n = 3. (B) IL-2 secretion from J-1E6 T cells co-cultured with either K562 A2/PPI or THP-1 cells. THP-1 cells fail to induce IL-2 secretion from J-1E6 cells even in the presence of exogenous $PPI_{15-24}$. Representative of n = 2 independent experiments. Statistical differences for (B) were determined by one-way ANOVA followed by Tukey's multiple comparison test. ns = non-significant, **** p < 0.0001.

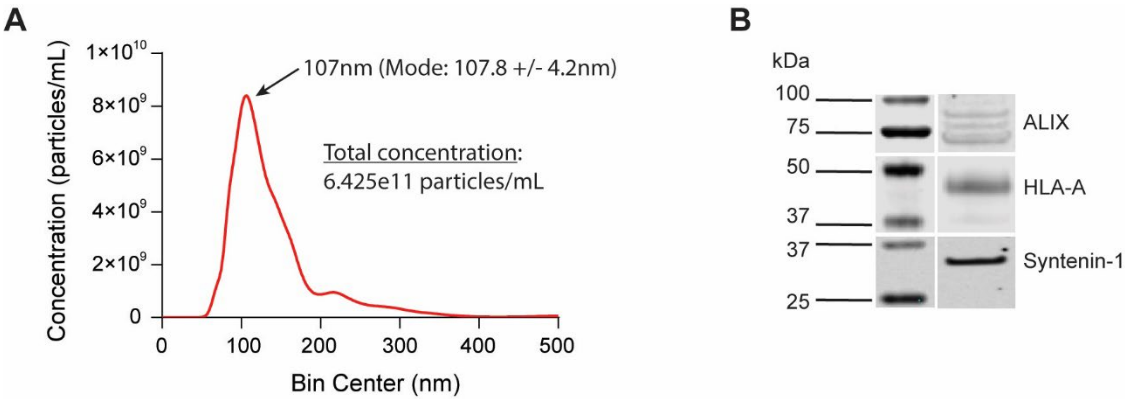


**Supplementary Figure 3**. Characterization of THP-1 EVs. (A) NTA showing representative particle concentration and size distribution of EVs derived from THP-1 cells. (B) Western blot analysis of exosome isolates demonstrating the presence of exosomal markers. Representative of n = 4 independent experiments.

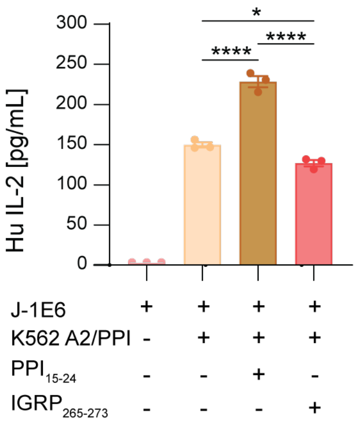


**Supplementary Figure 4.** Confirming antigen-driven activation of J-1E6 T cells in co-cultures. J-1E6 cells cultured with K562 A2/PPI cells and exogenous $PPI_{15\text{-}24}$ have increased IL-2 secretion compared to cultures without exogenous peptide, whereas culture with $IGRP_{265\text{-}273}$ peptide does not increase IL-2 secretion. Statistical differences for were determined by one-way ANOVA followed by Tukey's multiple comparisons test. * $p < 0.05$, **** $p < 0.0001$.

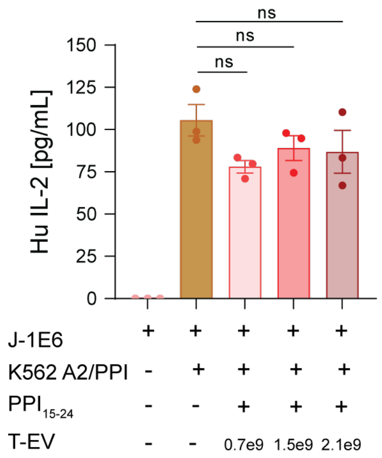


**Supplementary Figure 5.** Effects of THP-1 EVs on J-1E6 T cell activation. IL-2 secretion from J-1E6 T cells after co-culture with K562 A2/PPI cells and increasing amounts of T-EVs, showing no significant change in T cell activation with increasing amounts of EVs. n = 3. Statistical differences were determined by one-way ANOVA followed by Tukey's multiple comparison test. ns = non-significant.

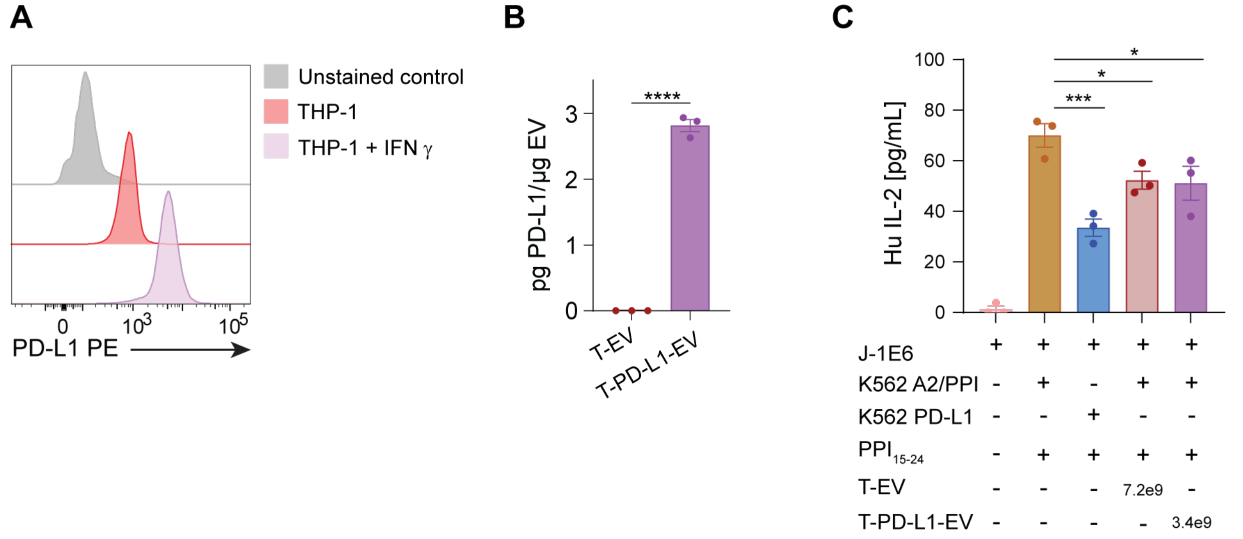


**Supplementary Figure 6.** Driving PD-L1 expression in THP-1 cells and EVs, and effects on J-1E6 T cell activation. (A) Surface expression of PD-L1 in THP-1 cells with or without IFN-γ treatment. n = 3. (B) ELISA of PD-L1 on EVs from THP-1 cells with or without IFN-γ treatment compared to total EV protein content. n = 3. (C) IL-2 secretion from J-1E6 T cells after co-culture with K562 cells and THP-1 EVs with or without PD-L1, showing slightly decreased T cell activation with excessive amounts of T-EVs or moderate amounts of T-PD-L1-EVs. Statistical differences for (B) were determined by an unpaired, two-tailed t-test. Statistical differences for (C) were determined by one-way ANOVA followed by Tukey's multiple comparison test. * $p < 0.05$, *** $p < 0.001$, **** $p < 0.0001$.
